# Fast genetic mapping of complex traits in C. elegans using millions of individuals in bulk

**DOI:** 10.1101/428870

**Authors:** Alejandro Burga, Eyal Ben-David, Tzitziki Lemus Vergara, James Boocock, Leonid Kruglyak

## Abstract

Genetic studies of complex traits in animals have been hindered by the need to generate, maintain, and phenotype large panels of recombinant lines. We developed a new method, C. elegans eXtreme Quantitative Trait Locus (ceX-QTL) mapping, that overcomes this obstacle via bulk selection on millions of unique recombinant individuals. We used ceX-QTL to map a drug resistance locus with high resolution. We also mapped differences in gene expression in live worms and discovered a regulatory feedback loop that responds to changes in HSP-90 chaperone activity. Lastly, we used ceX-QTL to map loci that influence fitness and discovered that one such locus is caused by a deletion in a highly conserved chromatin reader in the N2 reference strain. ceX-QTL is fast, powerful and cost-effective, and will accelerate the study of complex traits in animals.

## Introduction

Most heritable traits have a complex genetic architecture. Quantitative Trait Locus (QTL) mapping has been pivotal in identifying loci underlying complex traits of medical, agricultural and evolutionary importance^1–6^. However, genetic studies of complex traits remain challenging, especially in multicellular organisms. QTL mapping usually relies on generating large panels of cross progeny that must be individually genotyped and phenotyped. The construction and maintenance of such panels is lengthy, laborious and costly, limiting the size of most studies and their statistical power to confidently detect and narrow the genomic position of loci.

An alternative to traditional QTL mapping is Bulked Segregant Analysis (BSA)^7^. In the original BSA approach, cross progeny are still generated and phenotyped individually, but then individuals that fall into the tails of the phenotypic distribution are pooled for genotyping in bulk, and allele frequencies in the pools are compared to identify QTLs^8,9^. Building on the foundation of BSA and similar approaches^10^, our laboratory developed eXtreme Quantitative Trait Locus (X-QTL) mapping in the budding yeast *S. cerevisiae*^11^. In X-QTL, generation of cross progeny, genotyping and phenotyping are all carried out in bulk, enabling the use of extremely large populations of segregants (>10^6^ individuals), with correspondingly high statistical power and mapping resolution (Fig 1A). Using X-QTL, we have successfully resolved the genetic architecture of numerous complex traits in yeast, such as natural variation in resistance to chemicals, mitochondrial function, and gene expression^11–14^. However, we lack an equivalent powerful, fast and cost-effective method that can scale up to millions of individuals in animals.

**Figure 1.**
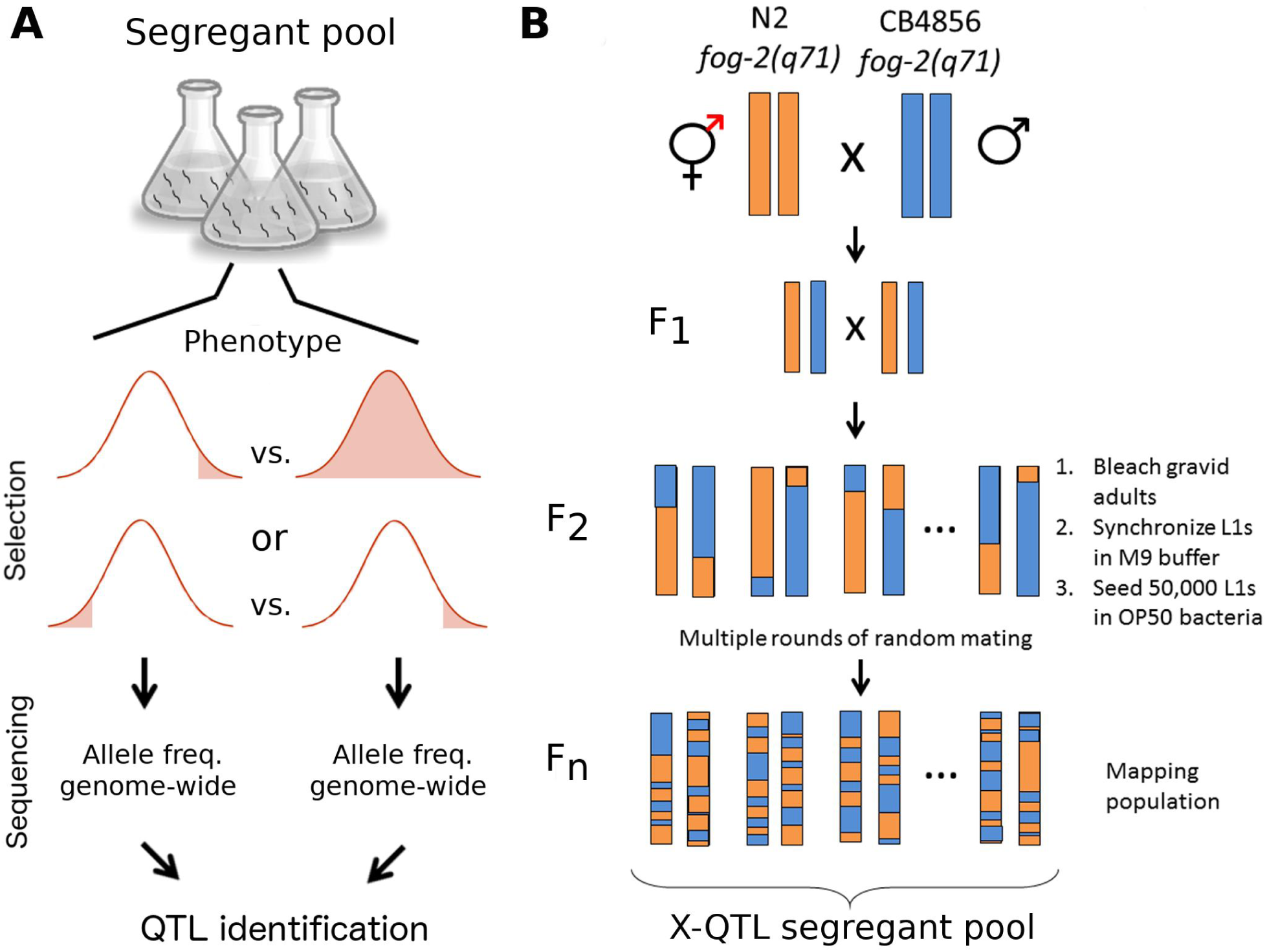
Implementing X-QTL in *C. elegans*. **(A)** The principle underlying X-QTL. A large population of recombinant individuals is selected based on a trait of interest. Two types of experiments are possible. Either one tail of the distribution is selected and contrasted with the overall population or two tails of the distribution are selected and contrasted with each other. Each group is then genotyped in bulk, and differences in allele frequency between these groups identify quantitative trait loci (QTL). **(B)** To generate millions of unique recombinant individuals in *C. elegans*, we used parental strains carrying the *fog-2(q71)* allele, which abolishes selfing. The resulting population can only reproduce by outcrossing. *C. elegans* experiences only one crossover event in each bivalent per meiosis, leading to an effective rate of half a recombination per chromosome. To increase the total number of recombination events and the resolution of *ce*X-QTL, we grew the segregant pool for multiple non-overlapping generations. Every generation, eggs were isolated by hypochlorite treatment from fully gravid adults. L1 larvae were synchronized by overnight starvation in M9 buffer and seeded on NGM plates (50,000 individuals).

Here, we extend the X-QTL approach to the nematode *C. elegans*. We show that *C. elegans* X-QTL (ceX-QTL) can be used to quickly map loci underlying differences in drug and stress resistance, gene expression, and fitness. Unexpectedly, our approach revealed that the *C. elegans* N2 reference strain is a knockout for *Y17G9B.8*, an ortholog of the highly conserved chromatin reader SGF29 (ref ^15^). The N2 *Y17G9B.8* allele is associated with transcriptional down-regulation of dozens of genes and decreased fitness. Both the yeast and human orthologs of *Y17G9B.8* specifically recognize methylated H3K4, mediate histone acetylation and enhance transcription, suggesting that the widely used N2 reference strain is not epigenetically wild type.

## Results

### Development of *C. elegans* X-QTL

X-QTL requires the generation of a large population of genetically unique segregants. Self-fertilization, the primary mode of reproduction of *C. elegans*, poses a challenge to implementing X-QTL because selfing individuals do not contribute to the genetic diversity of the segregant pool. To adapt the life cycle of *C. elegans* to X-QTL, we genetically abolished hermaphroditism using a *fog-2(q71)* mutation that ‘feminizes’ hermaphrodites by eliminating their sperm production^4^ (Fig. 1B). For simplicity, we will refer to these worms as females because they can only reproduce by crossing with males.

To generate the X-QTL segregant pool, we used two highly divergent *C. elegans* parental strains: the N2 reference strain (Bristol, UK) and the wild isolate CB4856 (Hawaii, USA). We constructed the CB4856 X-QTL parental strain by introgressing the *fog-2(q71)* allele into the Hawaiian background (Supplementary Fig. 1). We then crossed N2 *fog-2(q71)* females to CB4856 *fog-2(q71)* males and propagated a population of 50,000 segregants for 12 non-overlapping generations (Fig. 1B). We propagated the segregant population for multiple generations to increase the total number of recombination events per chromosome in the pool and, consequently, the mapping resolution^9,16^. Extensive simulations showed that this population size and number of generations provide sufficient genome-wide mapping power to detect loci explaining as little as 0.5% of phenotypic variance (Materials and Methods and Supplementary Fig. 2–4). A population of 50,000 is easy to maintain in the laboratory, and it can be quickly expanded to millions of individuals in a single generation because each female lays hundreds of eggs.

### Mapping natural genetic variation in drug resistance

Avermectins are a family of drugs widely used to treat parasitic worm infections and to fight insect pests. Thus, resistance to avermectins is a major health and agricultural problem^17^. We previously mapped a locus contributing to natural variation in Abamectin (Avermectin B1) resistance by studying the effect of this drug on locomotor activity in C. elegans^18^. Abamectin paralyses N2 at a faster rate than CB4856. To find the variant underlying this phenotypic difference, we originally performed QTL mapping using a large panel of 210 recombinant inbred advanced intercross lines (RIAILs). In each of these lines, sensitivity to Abamectin was determined by studying the frequency of body bends in liquid ^18^

In addition to affecting locomotor activity in adult worms, Abamectin is lethal to *C. elegans* larvae. We reasoned that resistance to Abamectin could be mapped in bulk by exposing a large number of N2 × CB4856 recombinant L1 larvae to a lethal dose of Abamectin and sequencing the surviving segregant pool. We treated four million F_12_ L1 recombinant larvae with 0.2 μg/mL of Abamectin in M9; only ~0.1% of the population survived this treatment (Fig. 2A). We extracted genomic DNA from the surviving population and from a control population that was exposed to DMSO alone. Finally, we estimated genome-wide allele frequency of 110,176 single-nucleotide variants (SNVs) using Illumina short-read sequencing and mapped QTLs by implementing a statistical framework previously developed for BSA^19^.

**Figure 2.**
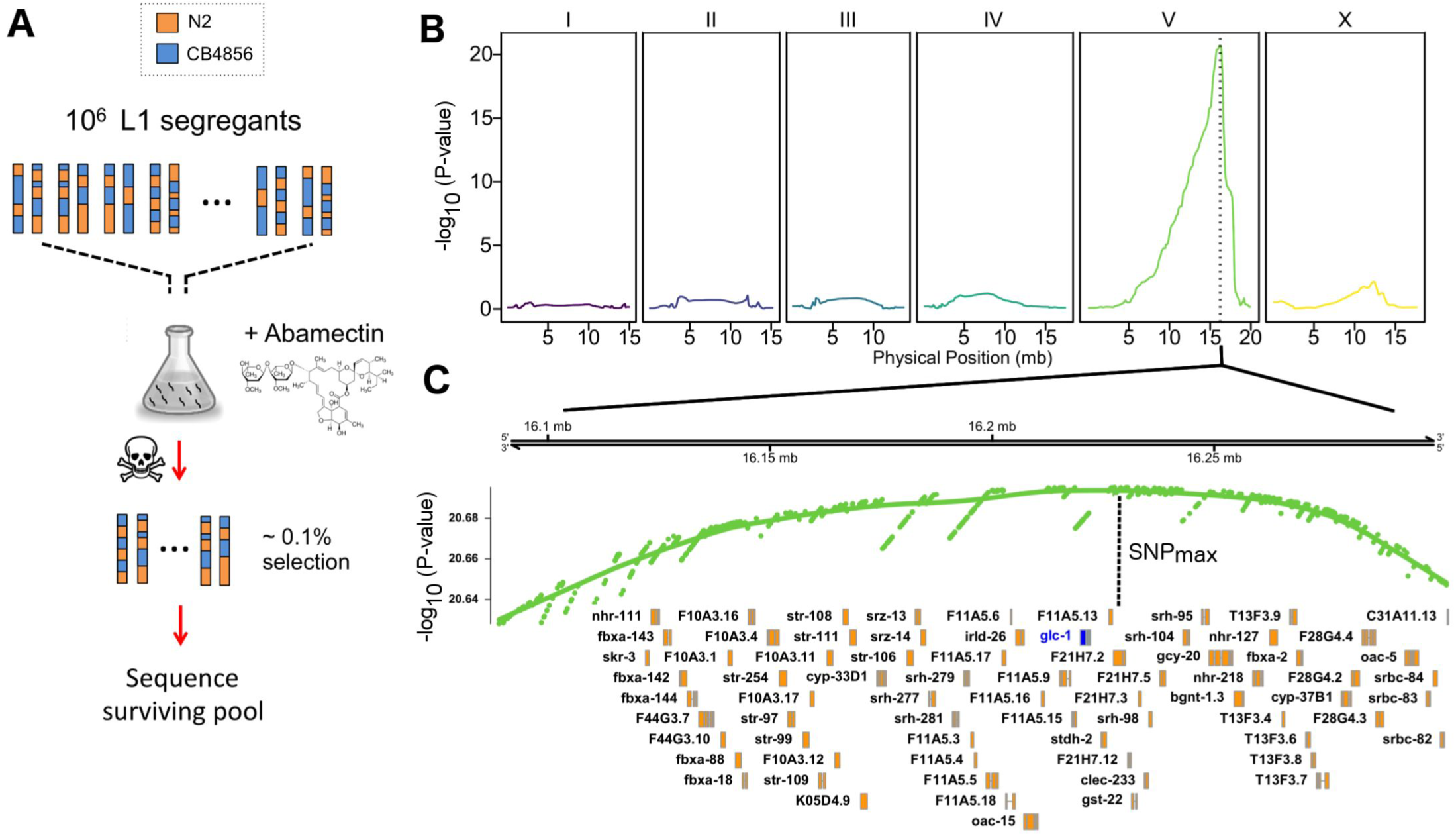
Mapping of natural resistance to Abamectin. **(A)** To map natural variation in Abamectin resistance using *ce*X-QTL, we treated four million F12 L1 recombinant larvae from a N2 × CB4856 cross with 0.2 μg/mL of Abamectin in M9 for 1 minute. Abamectin (Avermectin B1) is a natural fermenting product of the bacteria *Streptomyces avermitilis* and it acts as a nerve poison by binding invertebrate-specific glutamate-gated chloride channels. We then sequenced the surviving population (~0.1%) and a control population that was exposed only to DMSO (vector), and calculated genome-wide allele frequencies. **(B)** *ce*X-QTL identified a highly significant QTL peak on the right arm of Chr. V (p = 5.3 × 10^−22^). Dashed vertical line denotes the 95% confidence interval (CI): 16,115,957–16,276,907 Mb. **(C)** Focusing on the CI region, genes in orange. Dashed vertical line shows the SNV with lowest p-value in the region, which is located 3.7 kb away from *glc-1* (blue), the alpha-subunit of the glutamate-gated chloride channel. Natural variation in *glc-1* confers resistance to Abamectin.

*ce*X-QTL revealed a highly significant locus contributing to Abamectin resistance on Chr. V (Fig 2B, 95% confidence interval 16,115,957–16,276,907 Mb; *p =* 5.3 × 10^−22^), in the same region identified in the large RIAIL panel. The confidence interval obtained using *ce*X-QTL was smaller than the one obtained using the 210 RIAIL panel (224 kb for RIAIL panel^18^ and 160kb for *ce*X-QTL). The SNV with the most significant *p-value* was located only 3.7 kb away from the gene *glc-1* (Fig 2C). We previously showed that *glc-1*, which encodes the alpha subunit of a glutamate-gated chloride channel, is the causal gene underlying the QTL^18,20^.

In addition to drug resistance, *ce*X-QTL can also be used to map variation in any trait that can be selected for in bulk. For instance, we also subjected a population of 1.5 million L1 segregants to oxidative stress (0.5 mM H_2_O_2_) and uncovered 2 significant QTLs (Chr. II; p=1.60×10^−5^ and Chr. IV; p=3.07×10^−18^; Supplementary. Fig 5). These results illustrate the power of X-QTL in *C. elegans* to quickly guide the mapping of causal loci segregating in the wild in a single and fast experiment using a large mapping population.

### Coupling X-QTL and worm sorting

Our laboratory previously combined X-QTL and fluorescence activated cell sorting (FACS) to study the genetics of protein abundance in yeast ^14^. To develop an analogous approach in *C. elegans*, we coupled *ce*X-QTL to the Union Biometrica large-particle Biosorter, an instrument capable of viably sorting whole live worms. To demonstrate the power of this approach, we studied the transcriptional regulation of *C. elegans hsp-90 (daf-21)*, a highly conserved chaperone that is constitutively expressed throughout *C. elegans* development ^21^. To test whether *hsp-90* expression levels vary between isolates, we introgressed a single-copy *hsp-90p::GFP* transcriptional reporter from the N2 background into CB4856. We observed higher expression of this reporter throughout all developmental stages in the CB4856 background (2.6 fold upregulation in embryos; p=1.0×10^−4^ and 1.7 fold upregulation in adults; p=2.2×10^−6^; Supplementary Fig. 6). To map QTLs underlying this difference, we crossed the *hsp-90* reporter into our parental N2 *fog-2(q71)* and CB4856 *fog-2(q71)* X-QTL strains, and propagated a segregant population for 14 non-overlapping generations. We then measured the GFP fluorescence of ~60,000 F14 recombinant young adults and selected ~2,000 individuals from each of the two tails of the distribution (‘High’ and ‘Low’ GFP) (Fig. 3A, Supplementary Fig. 7). *ce*X-QTL analysis revealed a single highly significant locus on the left arm of Chr. V (*p* = 9.56 × 10^−69^, Fig. 3B). The ‘High’ GFP population was enriched for CB4856 alleles in the QTL region, as expected based on the parental phenotypes (Supplementary Fig. 7).

**Figure 3.**
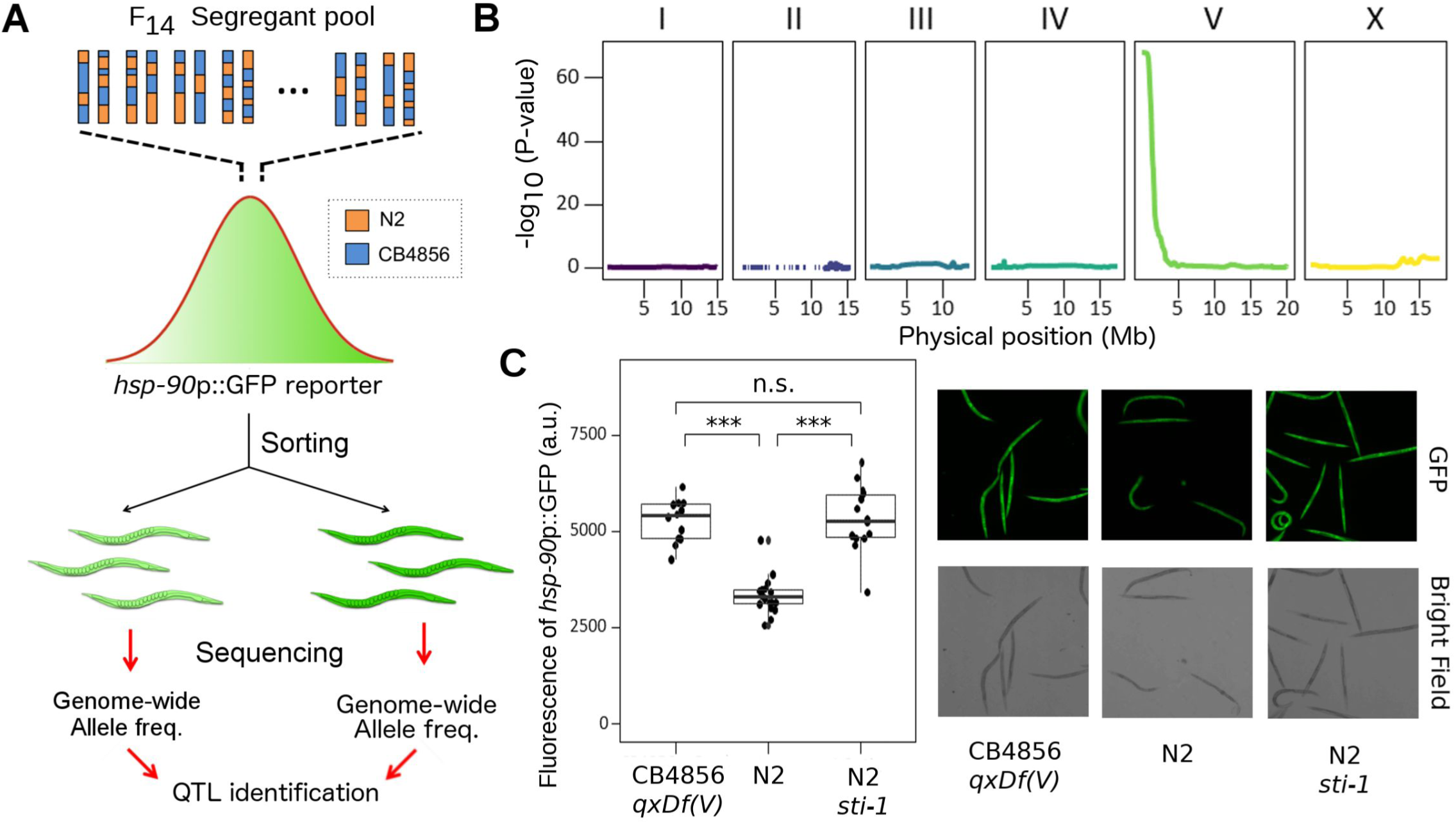
*ce*X-QTL identifies a regulator of *hsp-90* transcription. **(A)** To map variants underlying gene expression differences between isolates, we coupled *ce*X-QTL to a large particle biosorter. We crossed a transcriptional *hsp-90*::GFP reporter into our parental N2 *fog-2(q71)* and CB4856 *fog-2(q71) ce*X-QTL strains and propagated a segregant population for 14 generations. We then measured the GFP fluorescence of ~60,000 F_14_ recombinant young adults and selected ~2,000 individuals from the two tails of the distribution (‘High’ and ‘Low’ GFP). We sequenced both populations and calculated genome-wide allele frequencies. **(B)** *ce*X-QTL mapping identified a highly significant QTL peak in the left arm of Chr. V (p = 9.56 × 10^−69^). **(C)** A loss of function allele in the co-chaperone, *sti-1*, is sufficient to recapitulate the transcriptional upregulation of *hsp-90* observed in the parental CB4856 strain carrying *qxDf(V)*, a 267kb *de novo* deletion. Quantification of *hsp-90*::GFP reporter expression (left) and representative images (right). *Hsp-90* reporter expression was significantly higher in CB4856 *qxDf(V)* compared to N2 (1.58 fold upregulation, p = 7.9×10^−6^) and in N2 *sti-1(ok3354)* mutants compared to N2 (1.60 fold upregulation, p = 8.1×10^−6^). Hsp-90 reporter expression was not significantly different between CB4856 *qxDf(V)* and N2 *sti-1(ok3354)* (p =0.6)

Close inspection of the locus on Chr. V revealed a large 267kb deletion in the CB4856 *hsp-90p::GFP* strain (Chr. V:565,773–833,171 Mb) (Supplementary Fig. 6). This large deletion is not present in the original CB4856 parental strain, indicating that it was most likely acquired *de novo* during the introgression of the *hsp-90* transcriptional reporter into CB4856 and is not a natural polymorphism. Very little is known about the transcriptional regulation of *hsp-90*^22^ other than the role of *hsf-1*, the master regulator of the heat shock response. Therefore, we decided to further investigate the underlying causal mutation to gain insights into the regulation of this essential chaperone.

The Chr. V deletion encompassed 117 genes (Supplementary Table 1). We reasoned that the causal gene should be constitutively expressed during all developmental stages because the *hsp-90p::GFP* transcriptional reporter is upregulated in all tissues throughout the life of the worm. We leveraged gene expression data from the *C. elegans* modEncode project ^23^ and filtered our candidate list by keeping only those genes that were constitutively expressed during embryonic and larval development. This analysis reduced our list from 117 to 20 genes. We screened these 20 genes using RNAi on the parental *hsp-90p::GFP* N2 strain. RNAi silencing of only one gene, *sti-1*, caused upregulation of the *hsp-90* reporter (Supplementary Table 1). To confirm this finding, we studied a strain carrying the *sti-1(ok3354) allele*, a 336 bp deletion removing the last 90 amino acids of the protein. In agreement with our RNAi screen results, *sti-1* mutants showed upregulation of the *hsp-90p::GFP* reporter to a level indistinguishable from the CB4856 introgression strain (Fig. 3C). Together, these experiments indicate that *sti-1* loss of function causes transcriptional upregulation of *hsp-90*.

*C. elegans sti-1* is the ortholog of mammalian Hop and yeast Sti1, a co-chaperone that binds the chaperones Hsp90 and Hsp70^21,24^. Hop/Sti1 inhibits the ATPase activity of Hsp90 by acting as a non-competitive inhibitor and stabilizing the Hsp90 open conformation^25^. Since our *hsp-90p::GFP* transgene is exclusively reporting the transcriptional activity of *hsp-90* and not its protein abundance, our results suggest the existence of a novel positive feedback loop that responds to changes in HSP-90 chaperone activity by up-regulating its own transcription. Overall, our results illustrate how *ce*X-QTL in conjunction with a large particle biosorter can be used to study the genetics of gene expression in *C. elegans*. This strategy can be further extended to study natural variation in any trait amenable to sorting in living worms, including reporters of stress-response and lifespan^26^, mitochondrial activity^27^, maternal provisioning^28^, diet^29^, metabolism^30^, and neuronal activity^31^.

### *ce*X-QTL identifies loci influencing competitive fitness

Fitness, the measure of the reproductive success of an individual, is a complex genetic trait of fundamental importance to evolution. Without variation in fitness adaptation cannot occur. However, genome-wide mapping of genetic variants influencing fitness remains challenging, and has largely been limited to microorganisms. Throughout our experiments, we noticed that several genomic regions showed marked changes in allele frequencies that were shared by both our control segregant populations and those under selection. Although such baseline changes in allele frequencies do not hamper *ce*X-QTL mapping (Supplementary Fig. 9), we reasoned that they could reflect selective forces. To gain further insights into the origin of these allele frequency deviations, we sampled and sequenced earlier generations of the segregant pool, which were stored in frozen stocks.

We observed highly consistent deviations from the expected allele frequencies over the course of multiple generations (Fig. 4A). To exclude the possibility that these changes were the result of genetic drift, we independently generated and propagated two additional N2 × CB4856 segregant pools. Changes in allele frequencies were highly reproducible across biological repeats (Fig. 4A and Supplementary Fig. 10), suggesting that numerous genomic regions were most likely conferring fitness advantages.

**Figure 4.**
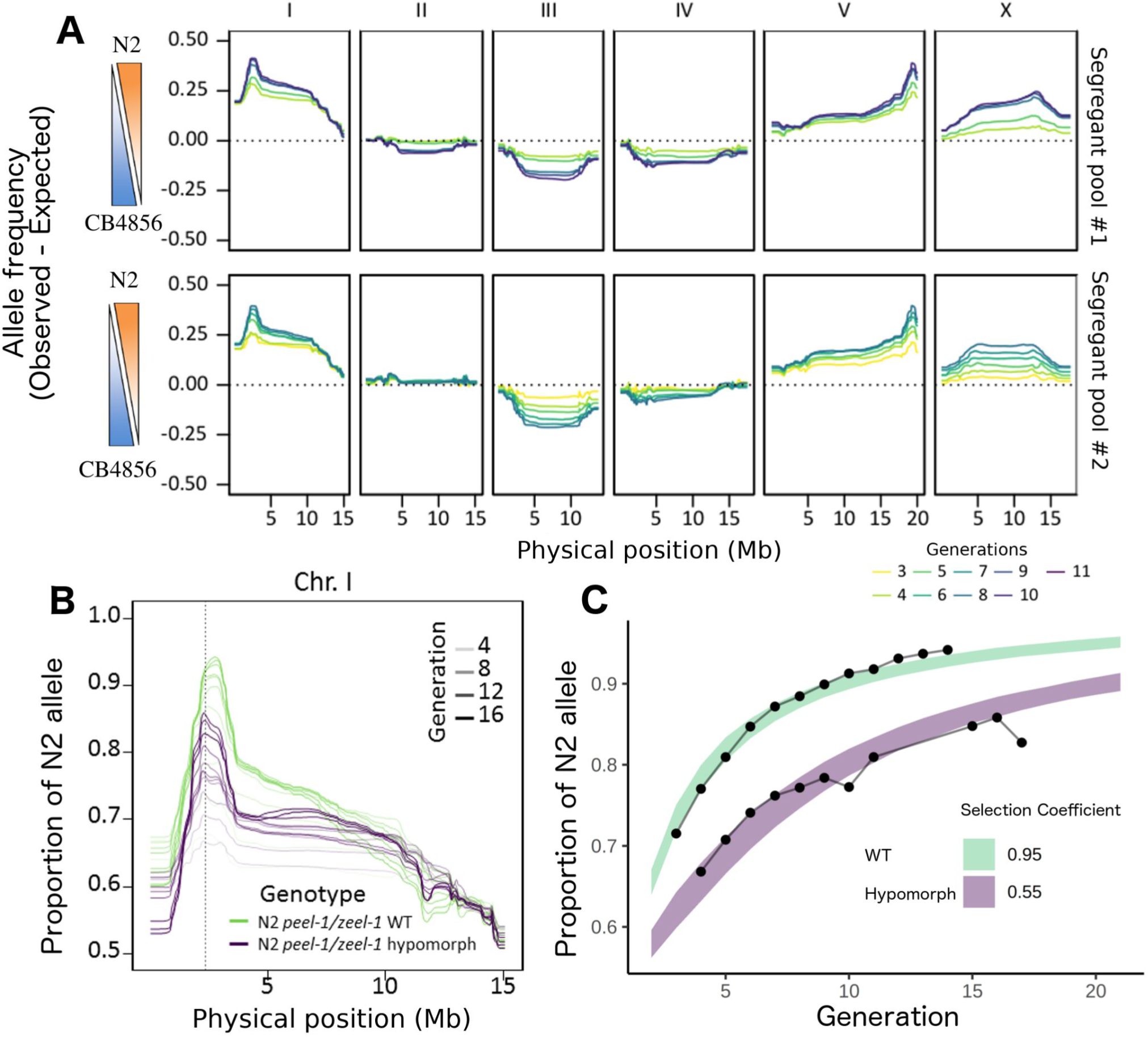
Temporal dynamics of ceX-QTL segregant pools reveal loci influencing competitive fitness. **(A)** Changes in allele frequencies across multiple generations (ranging from F_3_-F_11_) in two biological repeats of an N2 × CB4856 cross. Allele frequencies correspond to the difference between the observed and the expected values. **(B)** Driving activity of the *peel-1/zeel-1* selfish element in an N2 x CB4856 segregant population across multiple generations. N2 *peel-1/zeel-1* WT allele (green) and hypomorphic N2 *peel-1(ttTi12715)/zeel-1* allele (purple). The dashed vertical line denotes the location of the *peel-1/zeel-1* selfish element on Chr. I. **(C)** Estimation of selection coefficients from the observed changes in allele frequencies at the *peel-1/zeel-1* locus using simulations. The fitness of the poisoned homozygous individual lacking the selfish element is 1-s, where ‘s’ is the coefficient of selection. For a fully penetrant toxin-antidote element like *peel-1/zeel-1 allele, s*=1. The estimated ‘*s*’ for both WT and hypomorph alleles were 0.95 and 0.55, respectively. Bands are the 95% Confidence Intervals (CI) of simulations for WT (green) and hypomorph (purple).

One of the most extreme shifts in allele frequencies was observed on the left arm of Chr. I, where the *peel-1/zeel-1* selfish element is located^32,33^.The SNV with the largest deviation in allele frequency was located 40–100 kb away from the ~40 kb highly divergent structural variant spanning the *peel-1/zeel-1* element. This selfish element is composed of two tightly linked genes: *peel-1*, a sperm-delivered toxin and *zeel-1*, its zygotically expressed antidote. The N2 strain carries both genes, whereas, CB4856 lacks them. In crosses between heterozygous individuals, only the progeny that inherits at least one copy of the element survives, resulting in 25% embryonic lethality. We observed a progressive increase in the frequency of the N2 *peel-1/zeel-1* haplotype in the segregant pool, in agreement with its selfish ‘gene drive’ activity (Fig. 4A and B).

We wondered whether the observed changes in allele frequency of loci across generations could be used to quantify the relative strength of selection. To test this idea, we studied *peel-1(ttTi12715)*, a hypomorphic allele of the *peel-1* toxin carrying a transposon insertion^2^, and compared its fixation dynamics to those of the N2 wild-type allele. As expected, the drive activity of *peel-1(ttTi12715)* was not abolished, but it was reduced compared to that of the WT allele (Fig. 4B). To quantify this effect, we compared the observed results with simulations, allowing us to estimate the selection coefficient (‘s’). We found that selection was much weaker for the hypomorph, illustrating the sensitivity of our assay (s_wt_=0.95, s_hypomorph_=0.55; Fig 4C). Thus, our multigenerational *ce*X-QTL segregant approach is not only effective in mapping QTLs but it can also be used to quantify the relative strength of selection on loci influencing fitness.

A large peak on the left arm of Chr. X includes the neuropeptide receptor *npr-1*^34^. N2 carries a gain-of-function dominant mutation in *npr-1* that increases fecundity^35^. Replacement of the N2 *npr-1* allele with its CB4856 counterpart abolished the fitness peak on the right arm of Chr. X, thus confirming that *npr-1* was driving this signal (Supplementary Fig.11). In addition to known variants that contribute to fitness, we also uncovered several novel loci. For example, we found that almost the entirety of CB4856 Chr. III was selected over its N2 counterpart. This is particularly surprising because, in contrast to CB4856, the N2 strain has been selected for growth in the lab for over 50 years. We hypothesized that *plg-1* could underlie the strong selection in favor of CB4856 Chr. III due to male-male competition. *plg-1* encodes a mucin-like gene that is required in *C. elegans* to form a copulatory plug. In N2, this gene is disrupted by a transposon insertion, while it is functional in CB4856 and many other wild isolates^36^. However, reintroducing a functional *plg-1* allele into the N2 background did not affect the selection in favor of the CB4856 Chr. III, indicating that other unknown variants underlie this difference in fitness (Supplementary Fig. 11).

To further evaluate the reproducibility of the fitness peaks detected in our segregant pools, we studied a CB4856 *fog-2(kah89)* knock-in strain generated using CRISPR/Cas9 (Supplementary Fig. 11). We crossed N2 *fog-2(q71)* females to CB4856 *fog-2(kah89)* males and propagated the segregant pool for 17 generations. Notably, with the exception of a peak on the right end of Chr. V and a secondary peak in Chr. X, we could reproduce all the major fitness effect loci that we observed using the CB4856 *fog-2(q71)* introgression strain (Supplementary Fig. 10 and 11). The signal on the right end of Chr. V is most likely driven by a *de novo* mutation in close linkage with the *fog-2(q71)* introgression. These results show that direct gene editing by CRISPR/Cas9 is an effective method to generate parental ceX-QTL strains, and that it avoids effects that can arise from *de novo* mutations introduced during allele introgression.

### A highly conserved chromatin reader, *Y17G9B.8*, is disrupted in the N2 reference strain and decreases its fitness

Our data also revealed loci with antagonistic effects on fitness residing on the same chromosome. The first generations of our segregant pools showed weak but consistent selection in favor of CB4856 alleles on Chr. IV. However, by generation ten, it became apparent that the right arm of Chr. IV was being selected in favor of N2 (Fig. 4). We hypothesized that this selection pattern could emerge if variants with opposite effect on fitness were in linkage. To further examine this possibility, we propagated a ceX-QTL segregant pool for a total of 27 generations to accumulate more recombination events between the two fitness loci. Changes in allele frequencies in these advanced generations strongly suggested the presence of two independent loci on Chr. IV with antagonistic effects on fitness, with the left arm being selected in favor of CB4856 and the right arm in favor of N2 (Figure 5B and Supplementary Fig. 11).

**Figure 5.**
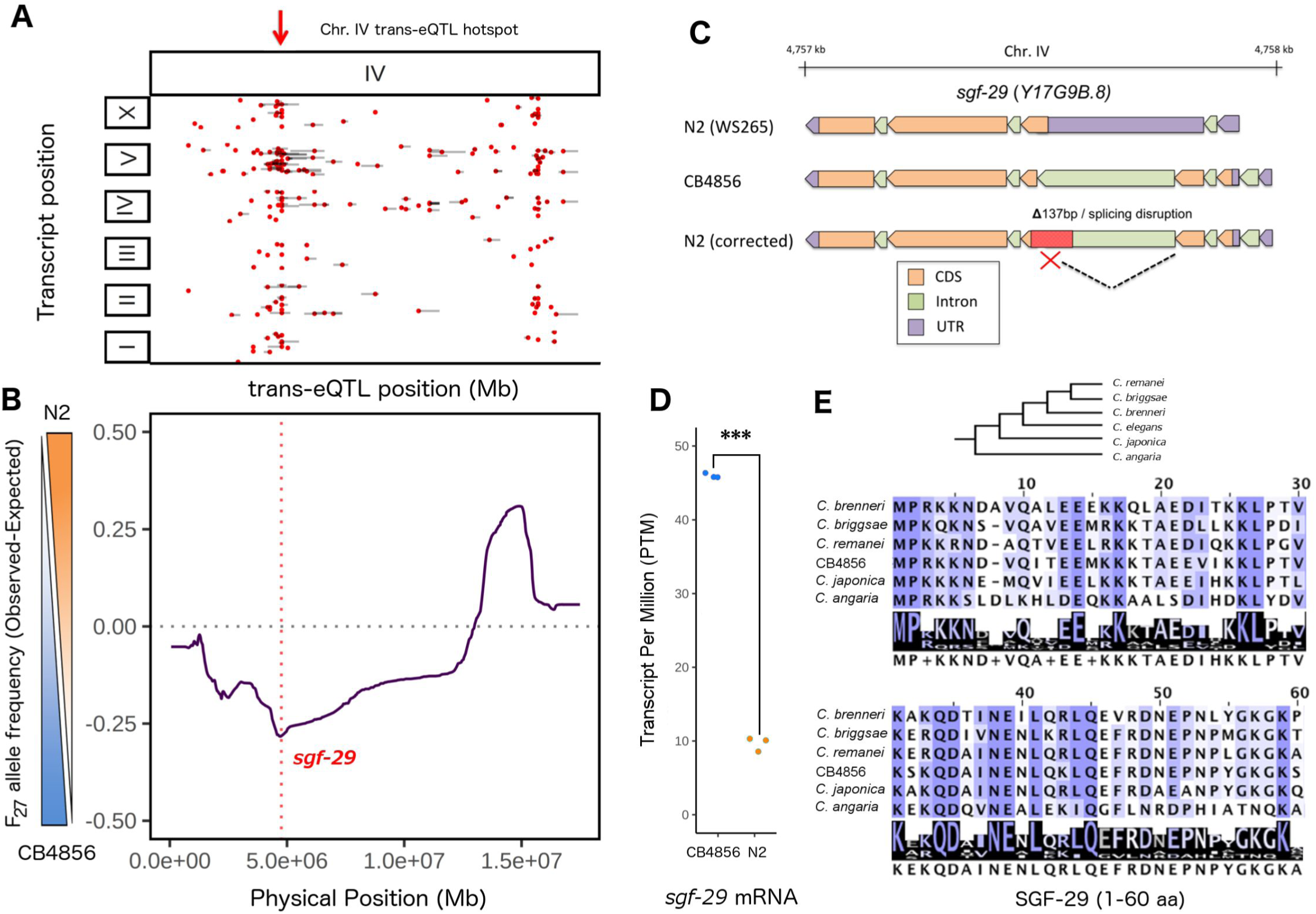
An eQTL hotspot in Chr. IV confers a fitness advantage to CB4856. **(A)** Position of distant eQTLs (red circles) mapping to Chr. IV in a N2 × CB4856 cross^37^, for simplicity local eQTLs are not shown. Gray bars represent 1-LOD drop confidence intervals. Position of the eQTL hotspot on the left end of Chr. IV is highlighted with a red arrow. This hotspot associates with the expression of 47 genes located across the *C. elegans* genome. **(B)** Allele frequencies of a *ce*X-QTL segregant pool expanded for 27 generations. Values correspond to the difference between the observed and the expected (0.5) in the absence of any selective force. The position in the genome of the gene *Y17G9B.8* is highlighted with a red dashed vertical line. **(C)** *Y17G9B.8* gene model. The current annotation of N2 *Y17G9B.8* on Wormbase WS265 (top), annotation of CB4856 *Y17G9B.8* based on RNA-seq expression data and evolutionary conservation (middle), and our corrected N2 annotation (bottom). A 137 bp deletion removes part of the third intron and 4 exon of *Y17G9B.8* in N2 causing intron retention. **(D)** RNA-seq quantification of *Y17G9B.8* transcript in the N2 and CB4856 parental strains (p=1.68×10^−6^, n = 3 for each strain) **(E)** Protein alignment of the first two coding exons of *Y17G9B.8*. These two exons were wrongly catalogued as a 5’ UTR in N2 WS265 but are highly conserved among *Caenorhabditis* at the protein level.

Our laboratory has previously studied the genetics of gene expression in a large panel of N2 × CB4856 recombinant inbred lines^37^. Comparing our *ce*X-QTL data with known expression QTLs (eQTLs) mapped in that study revealed that the fitness locus on the left arm of Chr. IV overlapped with an eQTL hotspot. (Fig. 5A and 5B). *Y17G9B.8*, an ortholog of the highly conserved chromatin reader SGF29, was originally regarded as a strong candidate gene for this eQTL hotspot because its own transcript abundance was linked to the locus, and because of its role in chromatin regulation^37^. Furthermore, a later study reported that the coding region of *Y17G9B.8* was disrupted by a large insertion in CB4856 compared to N2 (ref ^38^), pointing to the likely causal variant. But if *Y17G9B.8* is disrupted in CB4856, why would the *Y17G9B.8* CB4856 allele confer a selective advantage over N2 in our segregant pool? Re-annotation of this gene using short read RNA-seq data from N2 and CB4856 (ref ^39^), combined with evolutionary conservation analyses, revealed that, unexpectedly, *Y17G9B.8* is not disrupted in CB4856 as originally hypothesized^38^ (Fig. 5C and 5E). Instead, the putative insertion in CB4856 is in fact a 127 bp deletion that removes part of the third intron and the fourth exon of *Y17G9B.8* in N2, leading to intron retention predicted to disrupt translation (Fig 5C). In agreement with this, none of the 10 proteomics studies indexed in the pax-db database (http://pax-db.org) detected the predicted Y17G9B.8 peptide in N2, even though the transcript is consistently detected during all developmental stages^23^. Moreover, in agreement with previous microarray experiments, reanalysis of a published RNA-seq dataset^39^ showed that *Y17G9B.8* mRNA expression is significantly lower in N2 than in CB4856 (p=1.68×10^−6^, Fig 5D). Finally, conservation analysis revealed that the coding exons wrongly annotated as part of the 5’UTR in N2 (WS265) are highly conserved across *Caenorhabditis* nematodes (Fig 5E). Together, these results indicate that the N2 reference strain carries a null allele of *Y17G9B.8*. Among 152 *C. elegans* wild isolates that represent unique isotypes in the Caenorhabditis Natural Diversity Resource (CeNDR)^40^, the N2 *Y17G9B.8* deletion variant is only found in one other strain, LSJ1. N2 and LSJ1 are closely related lab-domesticated strains that diverged from a common ancestor in 1963^41^.

Laboratory-derived alleles in three genes—*npr-1, glb-5*, and *nath-10*—are known to increase the fitness of the N2 strain^2,35,42^. However, none of these variants are shared between N2 and LSJ1. Thus, the *Y17G9B.8* deletion allele is either one of the first genetic variants that arose during *C. elegans* domestication or a rare natural variant present in the original Bristol isolate, the ancestor of N2 and LSJ1. *Y17G9B.8* is an ortholog of human SGF29, a highly conserved member of the SAGA complex. Both yeast and human SGF29 have been shown to bind H3K4me2/3 marks and enhance transcription by mediating histone H3 acetylation^15^. Consistent with this role, the N2 allele was associated with decreased expression of 44 of the 47 target genes of the eQTL^37^. Overall, our results suggest that the reference N2 strain, used by the vast majority of worm researchers, may not necessarily reflect the common epigenetic state of *C. elegans*.

## Discussion

We have developed a novel method in *C. elegans* to quickly and cost-effectively dissect complex traits using millions of animals in bulk. Our approach can be readily adapted to other selfing nematodes. Furthermore, our bulk selection and genotyping approach will greatly facilitate studies of genetic variation in outcrossing nematodes^43^, where abolishing hermaphroditism is not required and inbreeding depression has hindered the generation of panels of inbred lines^44^. Our method offers many advantages over available mapping approaches. For any trait of interest that is amenable to bulk selection, it dramatically reduces the time and work required for genetic mapping compared to using large panels of RILs and wild isolates^2^. Moreover, it provides an internal control for phenotypic assays because all individuals are grown and selected in a homogeneous environment. Lastly, it can be easily expanded to study different genetic backgrounds without the need to construct, maintain, genotype, and phenotype large panels of recombinant lines. Once a *ce*X-QTL segregant pool has been generated, it can be used repeatedly to map different traits, as well as frozen for future use.

Our results also demonstrate the utility of *ce*X-QTL segregant pools to study selection and experimental evolution in *Caenorhabditis*^45^. In the competitive environment of our segregant pool, we have identified various loci influencing fitness, including an eQTL hotspot that originates from a deleterious mutation in a highly conserved chromatin reader in the reference N2 strain. The generation of *ce*X-QTL segregant pools is not restricted to two parental genotypes, thus allowing the study of multiple allelic variants in a single experiment. Currently, the main factor limiting the resolution of *ce*X-QTL and other mapping approaches is the highly nonuniform pattern of genetic recombination in *C. elegans* ^46^. Thus, we foresee that future implementations of *ce*X-QTL could greatly benefit from either targeting recombination to specific loci using CRISPR/Cas9^47^ or manipulating the endogenous recombination machinery of *C. elegans*^48^.

## Acknowledgements

We thank members of the Kruglyak lab for their comments. We thank Lijiang Long and Patrick T. McGrath for kindly sharing the CB4856 *fog-2(kah89)* strain. Funding was provided by the Howard Hughes Medical Institute and NIH grant R01 HG004321 (L.K.). A.B. is supported by the Jane Coffin Childs Memorial Fund for Medical Research. E.B. is supported by a Gruss-Lipper postdoctoral fellowship from the EGL foundation.

## Material and Methods

### Worm strains and growth conditions

*C. elegans* was grown using standard methods at 20°C^49^. Worms were fed with the *E. coli* OP50 on modified nematode growth medium (NGM), containing 1% agar and 0.7% agarose to prevent burrowing of CB4856. A detailed list of all the strains used in this study can be found in Supplementary Table 2. Some strains were provided by the CGC, which is funded by NIH Office of Research Infrastructure Programs (P40 OD010440). To generate the CB4856 *fog-2(q71)* strain, we introgressed the *fog-2* allele into CB4856 by performing nine rounds of backcross and selection for the feminization phenotype using CB4856 males. To confirm the introgression, we sequenced the resulting strain using Illumina short-read sequencing. The CB4856 *fog-2(q71)* strain carried only CB5856 variants with the sole exception of a ~ 1 Mb region in right arm of Chr. V where the *fog-2(q71)* allele is located (Supplementary Figure 1).

### Generation of *ce*X-QTL segregant populations

We transferred ~150 N2 *fog-2(q71)* virgin L4 hermaphrodites and ~150 CB4856 *fog-2(q71)* males into a single 5 cm. NGM plate. 24 hours later, worms and eggs were washed off from the plate and eggs were collected by hypochlorite treatment. The F_1_ generation was synchronized as L1 by starvation overnight in M9 buffer and seeded in three 15cm plates. NGM plates. After three days, once all the ‘females’ were fully gravid, eggs were once again isolated by hypochlorite treatment and L1s synchronized overnight in M9. We repeated this cycle for multiple generations, seeding 50,000 L1s every generation (~2,500 L1s per NGM plate) and freezing the rest of the population for long term storage. Due to the short life cycle of *C. elegans*, it only takes ~1 month to propagate the population for ten generations. Typically, we recovered a total of ~500,000 L1s every cycle. One generation before a *ce*X-QTL experiment, we further expanded the segregant pool by seeding >250,000 L1s in NGM plates. This expansion guaranteed the recovery of over a million L1 segregants the next generation readily available for mapping. If required, tens of millions of segregants can be easily obtained by carrying out two population expansion cycles in ~1 week. Importantly, a single *ce*X-QTL segregant pool can be used for multiple mapping experiments or it can be frozen for long term storage. We have successfully thawed and propagated glycerol stocks of segregant pools and used them for mapping experiments.

### Generation of variant list in CB4856

To assemble a list of variants between N2 and CB4856, we reanalyzed published Illumina sequencing of CB4856 ^40^ Reads were aligned to *C. elegans* reference build WBcel235 and variants were called using four different genotyping software: *platypus* ^50^, *varscan* ^51^, *Freebayes* ^52^ and the Genomic Analysis Toolkit (GATK) ^53^ We considered only SNVs identified by at least three methods. The entire process was automated using the *bcbio-nextgen* pipeline *(ver. 0.9.9) (https://bcbio-nextgen.readthedocs.io/)*. The analysis identified 227,228 SNVs. These SNVs were used to generate a custom reference for CB4856. We then filtered these SNVs further, by aligning reads from our CB4856 *fog-2(q71)* strain to both the reference N2 and our custom CB4856 genome build and retaining only SNVs in which over 90% of the reads supported the N2 allele in both alignments. We further excluded SNVs in the mitochondrial DNA. The final list of SNVs included 110,176 variants.

### Generation and processing of Illumina whole-genome sequencing

Genomic DNA was extracted using the DNeasy Blood & Tissue kit (Qiagen). Illumina sequencing libraries were prepared using the Nextera DNA Library Prep Kit (Illumina). We followed the standard protocol with the following exception: we performed agarose size-selection of the Nextera libraries, extracting a ~500 bp band. Libraries were sequenced on Illumina Miseq, Hiseq 2500, Hiseq 4000 and Hiseq X sequencers (see Supplementary Table 3 for a description of all sequencing runs). Reads were aligned to the WBcel235 genome. Alignment bam files were sorted and filtered of PCR duplicates using *sambamba* ^54^ Finally, allele counts in each SNV were calculated using the program *bam-readcount* (https://github.com/genome/bam-readcountj.

### Statistical analysis of *ce*X-QTL

We implemented a previously published statistical method developed by Magwene et al.^19^. For each SNV, allele counts are used to calculate a G statistic. To account for segregation distortion, we used a modified version of the statistic estimated in Magwene et al: 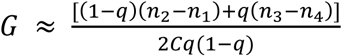, where n_1_ and n_3_ are counts of allele A and B in the high (or treatment) population, and n_2_ and n_4_ are counts of allele A and B in the low (or control) population; C is the depth of coverage at each SNV, and q is the baseline frequency of allele A (estimated using the control/low population). The raw G statistic was smoothed using a weighted average approach, where the smoothed G’ statistic for SNV s is given by 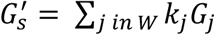, where *k* is the genetic distance between SNV *j* and *s*, transformed using the tri-cube kernel function 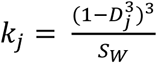; *W* is a window around *s*, so that only SNVs within the window are used to calculate the weighted average. We imposed a cutoff of 12.5cM (*W* = 25cM), the lower bound of the values suggested by Magwene et al. To calculate p-values corresponding to values of G’, the null distribution of G’ was estimated from the data using a robust fit to the log-normal distribution (as implemented by the *robust* R package). We’ve written an R package that implements the entire statistical pipeline, *xQTLstats*, and it is available on github (github.com/eyalbenda/xQTLstats).

### Simulations of *C. elegans* segregant populations

To determine the power of X-QTL in *C. elegans*, we first sought to accurately simulate the process of propagating a segregant population. We developed a simulation framework, *bulkPop*, where each individual is represented by two haplotype vectors. Mating is implemented in a straightforward way as a process whereby the haplotypes in each parent recombine, followed by random independent segregation of the recombined haplotypes to progeny. Reflecting the recombination rates in *C. elegans*, the probability of each chromosome to undergo a single recombination event was 0.5, and the location of the recombination was determined using a genetic map based on a recombinant inbred line panel between CB4856 and N2^46^. To simulate loci conferring a fitness advantage (“fitness loci”), we randomly select a subset of individuals from the population to produce progeny, and probability of an individual to be chosen to mate was weighted by the genotype in fitness loci. Lethality due to the *peel-1/zeel-1* element is a result of an interaction between the parental and the zygotic genotype, and this interaction was directly simulated to accurately predict the segregation distortion due to the element. Our framework can easily be extended to crosses in other strains or oganisms. It is implemented as an R package, *bulkPop*, and is available on github (github.com/eyalbenda/bulkpop).

### Simulating the power of *ce*X-QTL

We used *bulkPop* to propagate a large population for 10 non-overlapping generations. The starting F1 population was 1,000 worms, and the population was capped at 50,000 worms, with each mated female generating 10 progeny. After 10 generations, the population was expanded to 1 million worms 100 times, generating a large 100 million “pool” of segregants to use for simulations. To determine the effect of fitness loci on the power of *ce*X-QTL, fitness was modeled as affecting the probability of a male to participate in mating. All loci were modeled as driven by a single factor, and the strength of selection was chosen such that the segregation distortion in the simulated population was similar to the observed distortion in the X-QTL population across generations.

On the large populations, a *ce*X-QTL drug selection experiment (Figure 1A) was simulated directly by selecting a 5% survivor population, with a random subset of the large population selected as control. To select 5% survivors in a way that guaranteed that loci had a specified effect size (modeled as the variance explained by the locus *V_e_*), the following procedure was used:

a. the genotype of each individual in the causal locus was encoded as *g* = {0,1,2}.
b. For each individual, a random displacement factor *d* was simulated for each individual from a normal distribution, with *μ* = 0, and 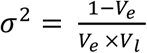, where *V_l_* is the variance of the vector of genotypes *g* in the causal locus.
c. The final score of each individual was *S = g + d*

On that score, a cutoff of 5% was imposed to select the survivors of drug selection. Allele counts were simulated based on the allele frequencies in each population using the binomial distribution and used as input for *xQTLstats*.

### Selection for Abamectin resistance

We propagated a N2 × CB4856 segregant pool for 12 generations in NGM plates. We incubated 4 million F_12_ L1 larvae tin 0.2 μg/mL Abamectin (Sigma-Aldrich) in M9. Abamectin was freshly dissolved from a 10 mg/mL stock in Dimethyl sulfoxide (DMSO) kept at −20C. L1 larvae were incubated in Abamectin for 1 minute and washed three times with 15 mL of M9 buffer. After the washing steps, L1 larvae were seeded on OP50 NGM plates at a density of ~200,000 larvae per plate. Approximately 0.1% of larvae survived this treatment and developed into adult females and males. Surviving adults were washed off from plates and pooled for DNA extraction. As a control, ~5,000 F_12_ larvae from the same population were exposed to an equivalent dose of the vector DMSO for 1 minute, seeded on OP50 NGM plates and collected for DNA extraction when the population developed into adults.

To determine the 95% confidence interval for the identified QTL, we simulated a drug selection study as described above, simulating a drug resistance variant explaining 10–25% of the variance.

### Selection for H_2_O_2_ resistance

We propagated a N2 × CB4856 segregant pool for 10 generations in NGM plates. We exposed 1.5 million F_10_ L1 larvae to 0.5M H_2_0_2_ (Sigma-Aldrich) in M9 buffer for 4 hours. Approximately 0.1% of larvae survived this treatment and developed into adult females and males. Larvae were washed three times with 15 mL of M9 buffer. After the washing steps, L1 larvae were seeded on 0P50 NGM plates at a density of ~200,000 larvae per plate. Approximately 0.1% of larvae survived this treatment and developed into adult females and males. As a control, ~5,000 F_12_ larvae from the same population were incubated in M9 for 4 hours, seeded on OP50 NGM plates and collected for DNA extraction collected for DNA extraction when the population developed into adults.

### Selection for hsp-90 transcriptional reporter expression levels

A single copy^55^ *hsp-90p::GFP::hsp-90* 3’UTR transgene reporter in the N2 genetic background (BCN1082^56^) was introgressed into the CB4856 genetic background by performing six rounds of backcross and selection. We then crossed the N2 and CB4856 *hsp-90p*::GFP reporter lines to their respective *ce*X-QTL parental strains carrying the *fog-2(q71)* mutation. The resultant strains QX2314 (N2 *fog-2(q71)* V; *hsp-90::GFP II*) and QX2307 (CB4856 *fog-2(q71)* V; *hsp-90::GFP* II) were used to generate a *ce*X-QTL segregant population. Fluorescence based sorting was carried out with a Large Particle Biosorter (Union Biometrica) equipped with a 250μm Fluidics and Optics Core Assembly (FOCA). Worms were grown on plates to young adult (2.5 days after seeding synchronized L1 larvae), and washed into 50ml conical tubes (target concentration 1 worm/μl). Worms were anaesthetized by adding Levamisole to 3mM final concentration.

### RNAi screening

To determine a list of candidates in the deletion on Chr. V, we reanalyzed RNA-seq data from the ModEncode project representing gene expression from large synchronized worm populations collected at different developmental stages^23^. Gene expression was quantified from raw sequencing reads using Kallisto^57^. We selected 20 genes that were expressing throughout life for RNAi screening. We used the Ahringer RNAi library (Source Biosciences). We blindly screened RNAi clones targeting 20 genes (Supplementary Table 1) in NGM plates supplemented with Ampicillin (100μg/ml) and IPTG (1mM). N2 worms carrying a hsp-90::GFP single copy reporter were scored for increased GFP fluorescence using a stereoscope equipped with a fluorescence lamp.

### Microscopy

Worms were transferred to a 3% Agarose pad and visualized using a Nikon Eclipse 90i microscope equipped with a Photometrics CoolSNAP HQ2 CCD camera.

### Testing candidate variants affecting fitness

We crossed the parental CB4856 *fog-2(q71)* strain to a modified N2 *fog-2(q71)* strain carrying a hypomorphic *peel-1/zeel-1(ttTi12715)* allele carrying a transposon insertion (Chr. I, introgressed from QX1430), a functional *plg-1* allele (Chr. III, introgressed from CB5203) and the WT (CB4856) allele of *npr-1* (Chr. X, introgressed from QX1430). All the alleles were verified in the final strain by PCR or Sanger sequencing.

### Estimation of selection coefficients

We used our *bulkPop* package to simulate the effect of the *peel-1/zeel-1* element on allele frequencies under 20 different values of selection coefficient, which was modeled as the penetrance of the element between 0 (no lethality) and 1 (full lethality). For each value, we simulated 50 populations of 10,000 worms for 20 generations. To estimate the observed selection for the *zeel-1/peel-1* element in the ceX-QTL populations, we took the maximum value *wild-type (wt)* All ceX-QTL experiments that were done with worms carrying the *wt* allele were averaged. We determined the best fit to the simulated population by finding the value minimizing the sum of the differences between the average of the simulations and the observed across all generations.

### Analysis of the fitness peak in Chr. IV

Microarray genotype and gene expression data for our published expression QTL data^37^ were acquired from the gene expression omnibus (GEO). To eliminate discrepancies in gene annotations, probe sequences were realigned to the WBcel235 transcriptome using BWA^58^.Only uniquely mapping probes were used. Expression probes that were present in less than 2/3 of the sample were removed. The genotype and expression matrices were normalized to have mean zero and variance one. To map eQTLs, we calculated the Pearson correlation between each probe and every genotype. Correlation coefficients were transformed to LOD scores using 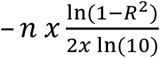. To assess significance and account for multiple testing, we permuted the sample identities 100 times and calculated the average number of transcripts with an identified eQTL at different LOD scores. We compared these results to the unpermuted LOD scores to estimate the false-discovery rate (FDR)^59^, and selected a cutoff corresponding to a rate of 5%. To compare the expression of *Y17G9B.8* in N2 and CB4856 we quantified gene expression in a previously published dataset using Kallisto^57^. To ensure that the estimates would not be skewed because of the deletion in N2, we used the N2 version of the gene as the reference (as available in the ensembl WBcel235 transcriptome). Differential expression analysis was performed using Sleuth^60^ and *p-value* for differential expression was calculated using the likelihood ratio test as implemented in Sleuth.

## Supplementary Figures

**Supplementary Fig. 1.**
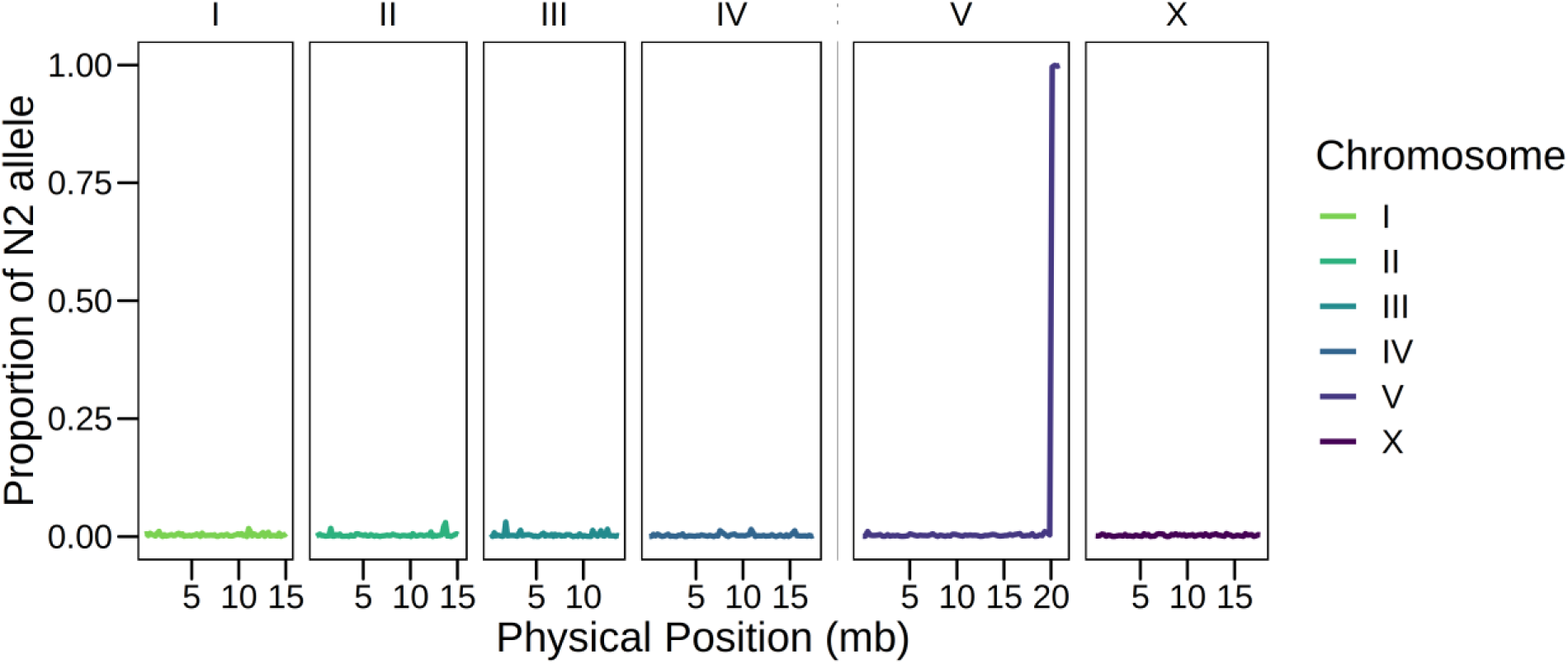
Verification of the CB4856 *fog-2(q71)* introgression strain. Genotyping of the introgression strain was performed using Illumina short read sequencing after 9 backcrosses into CB4856 parental strain. The resulting strain contains a ~1 Mb N2 region in the right end of Chr. V where the *fog-2* gene is located. The introgressed region is not considered for X-QTL mapping.

**Supplementary Fig. 2.**
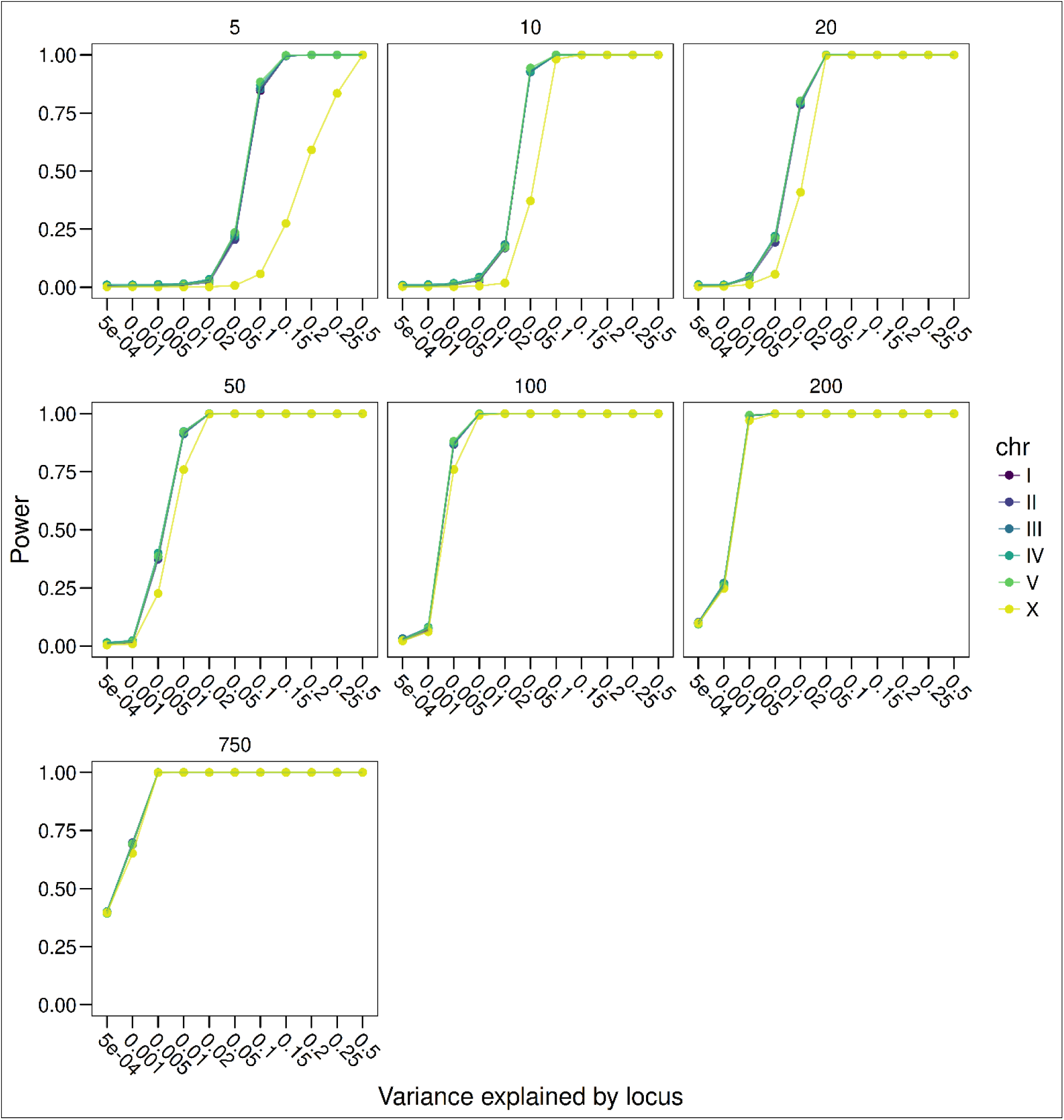
Worm X-QTL is a powerful method for mapping traits across a spectrum of effect sizes. An X-QTL for drug selection (survivors VS. background) was simulated. In each simulation, a population of 50,000 simulated F10 segregants were subjected to a selection with 5% survivors. Loci affecting survival were simulated with effect sizes ranging from 0.05% to 50% of variance explained. Allele counts were simulated from each population with sequencing depth ranging from 5x to 750x. 10 equally distanced loci were simulated across each chromosome. With coverage of 100x (~10GB of sequencing), X-QTL has power to detect loci explaining as little as 0.5% of the phenotypic variance.

**Supplementary Fig. 3.**
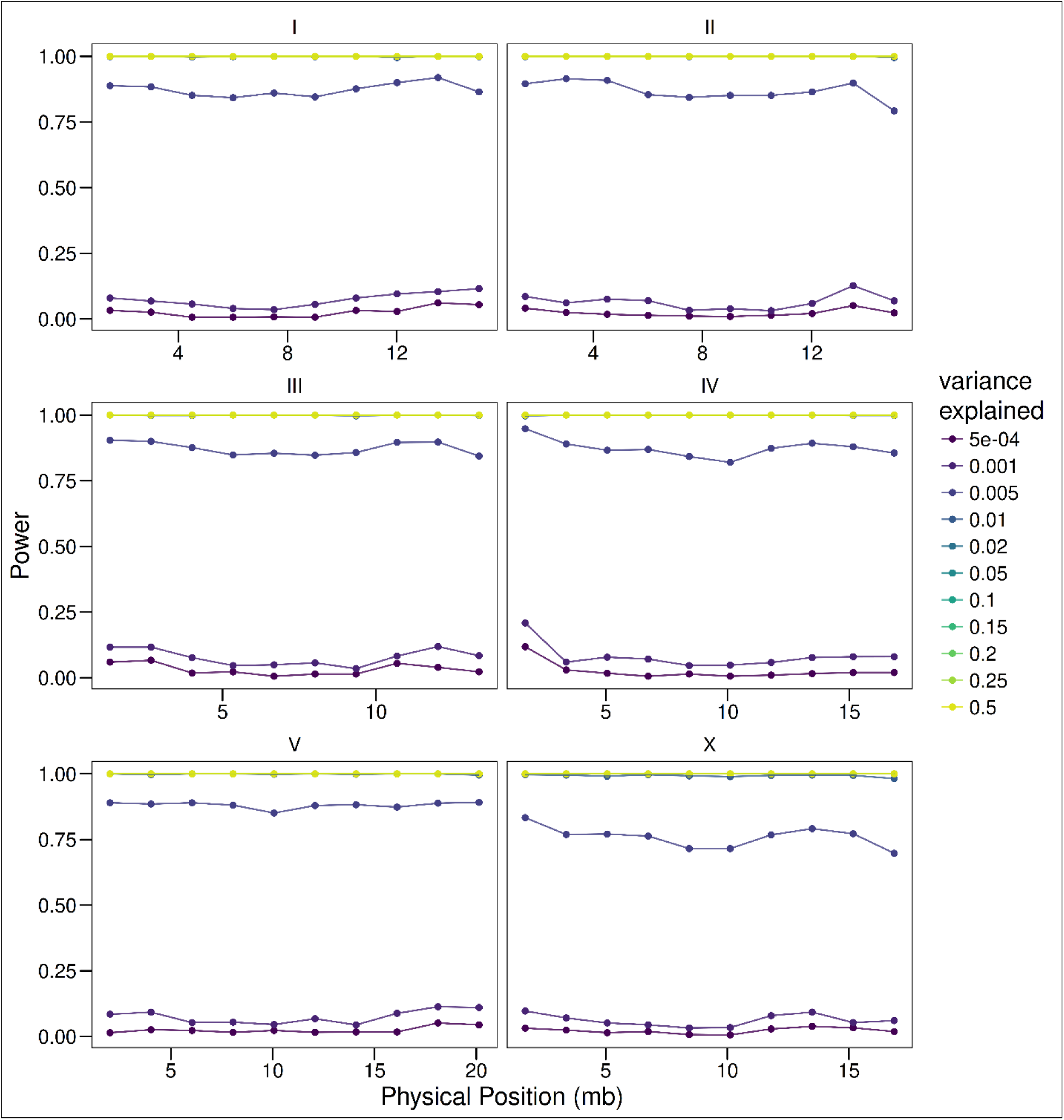
Power of X-QTL across chromosomes. Simulations were carried out as described in Supplementary Fig. 2. 10 loci were simulated at equal distances across each chromosome. With a sequencing depth of 100x (~10 GB of sequencing), X-QTL maintains the power to detect loci above 0.5% variance explained across the *C. elegans* genome.

**Supplementary Fig. 4.**
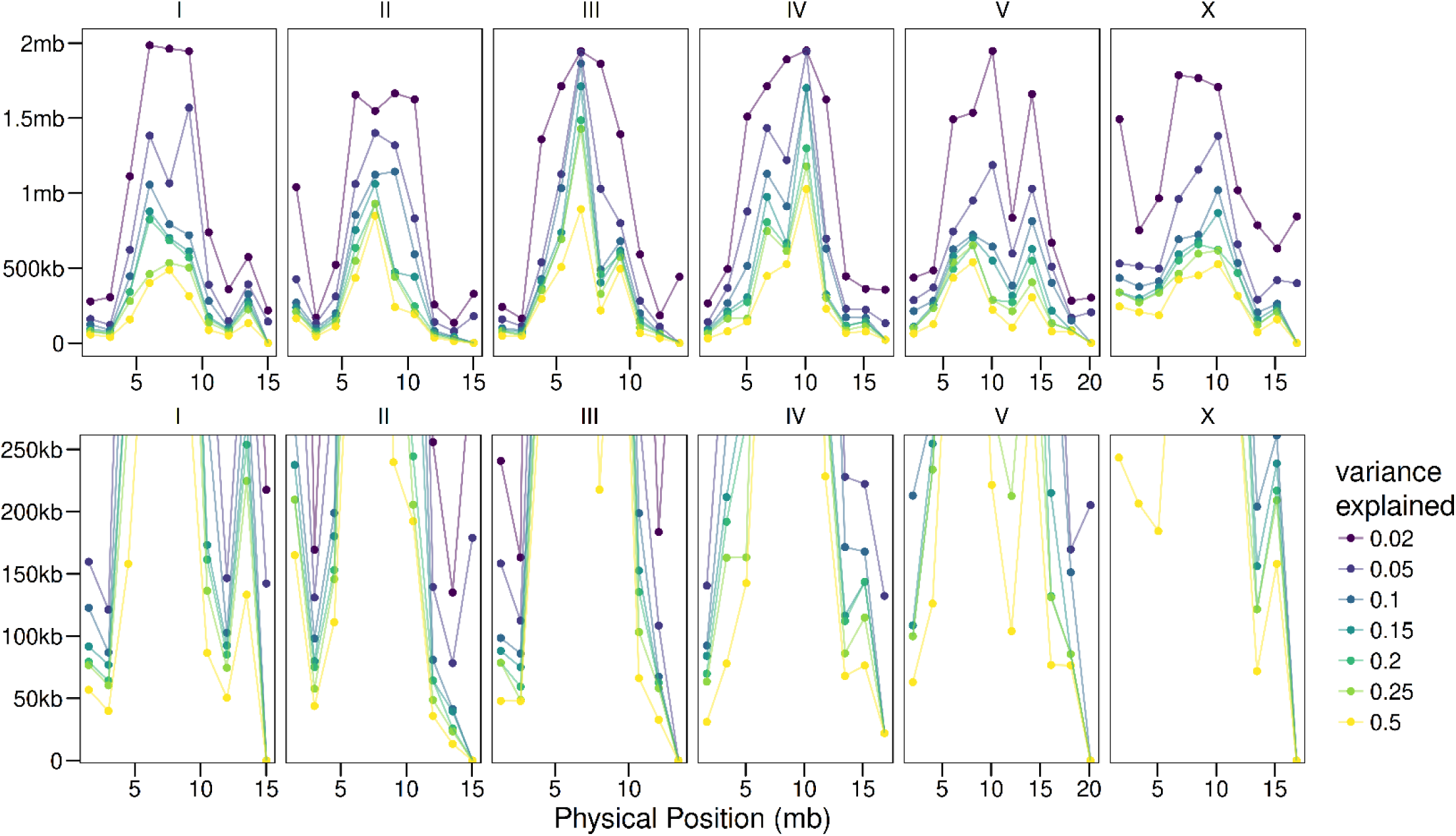
Resolution of X-QTL varies across the chromosome. Simulations were carried out as described in Supplementary Fig. 2, focusing on simulations with a sequencing depth of 100X. For each simulation, the peak position of the mapped X-QTL peak was taken, for a total of 1000 repeats per position. The resolution was defined as the range encompassing 95% of the peaks. As expected, the resolution varies widely across *C. elegans* chromosomes, due to the highly uneven recombination rates. Bottom panel focuses on a range from 50kb-250kb, while top panel shows the entire range.

**Supplementary Fig. 5.**
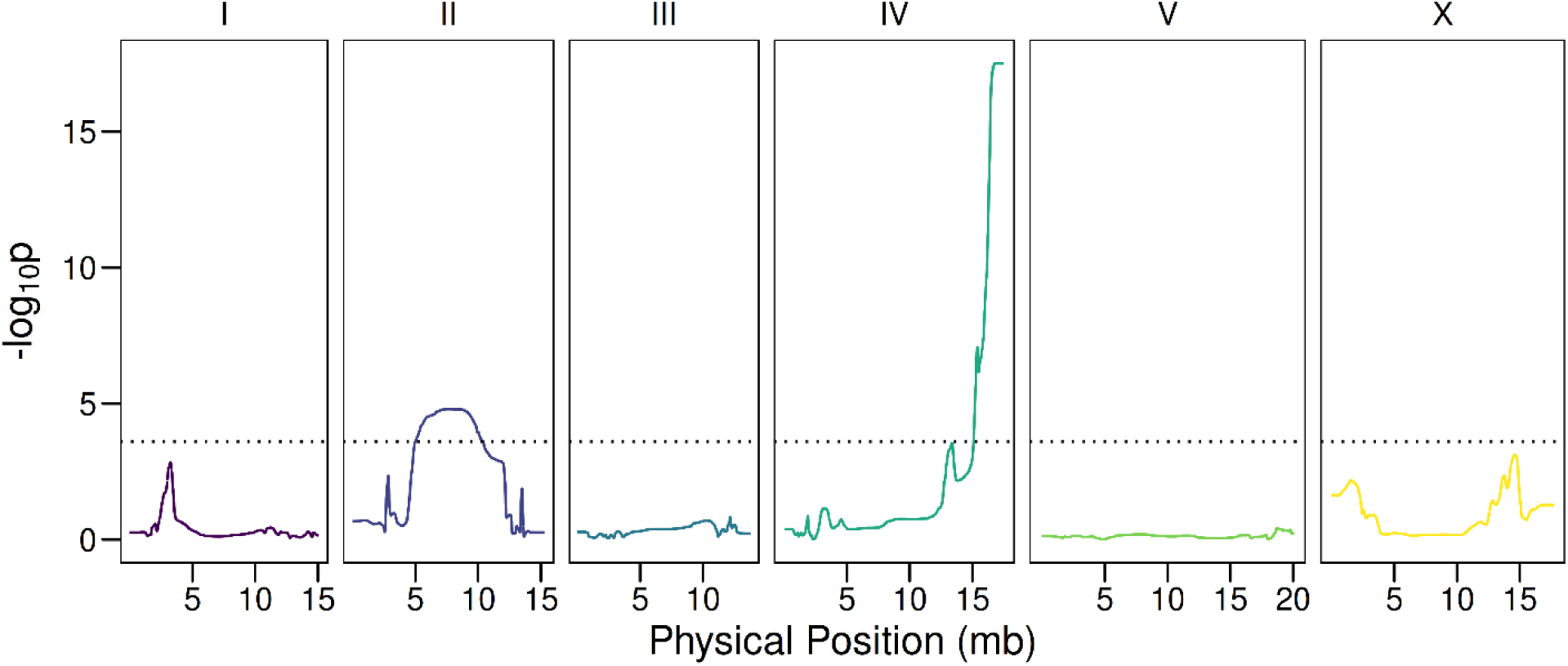
X-QTL mapping identifies loci conferring resistance to H_2_0_2_. Two significant loci were identified, on chromosomes II (p=1.60×10^−5^) and IV (p=3.07×10^−18^)

**Supplementary Fig. 6.**
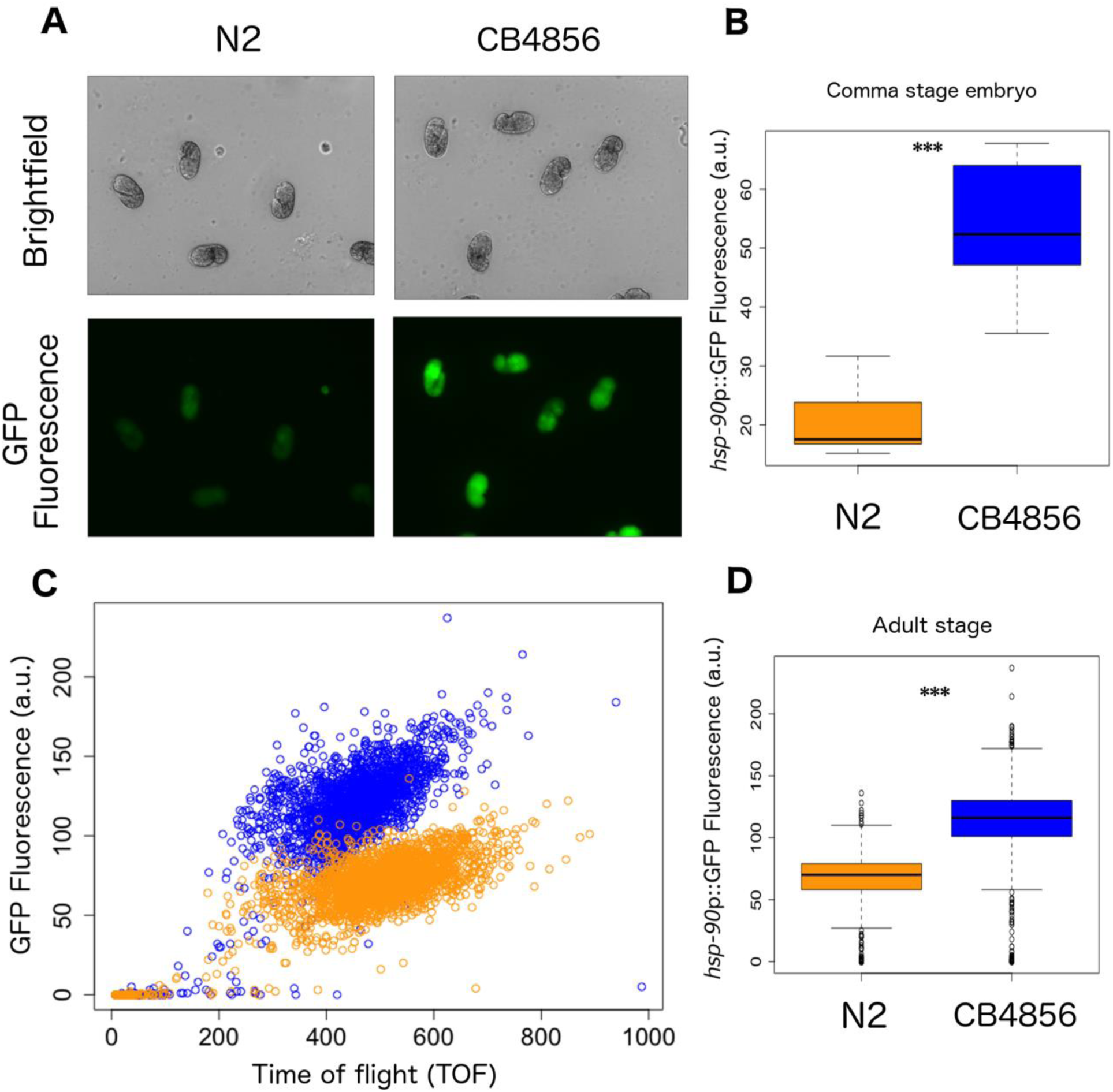
*Hsp-90::GFP* reporter expression levels are upregulated in the CB4856 background. **(A)** Representative images of *hsp-90::GFP* comma stage embryos in both N2 and CB4856 background. **(B)** Quantification of GFP fluorescence. *hsp-90::GFP* is upregulated in CB4856 (2.6 fold upregulation; p=1.0×10^−4^, n=7 embryos). (C) Quantification of GFP fluorescence in adult N2 and CB4856 *hsp-90::GFP* expressing worms using a COPAS Biosorter. **(D)** *hsp-90::GFP* is upregulated in CB4856 (1.7 fold upregulation; p=2.2×10^−6^, n= ~2,500 individuals)

**Supplementary Fig. 7.**
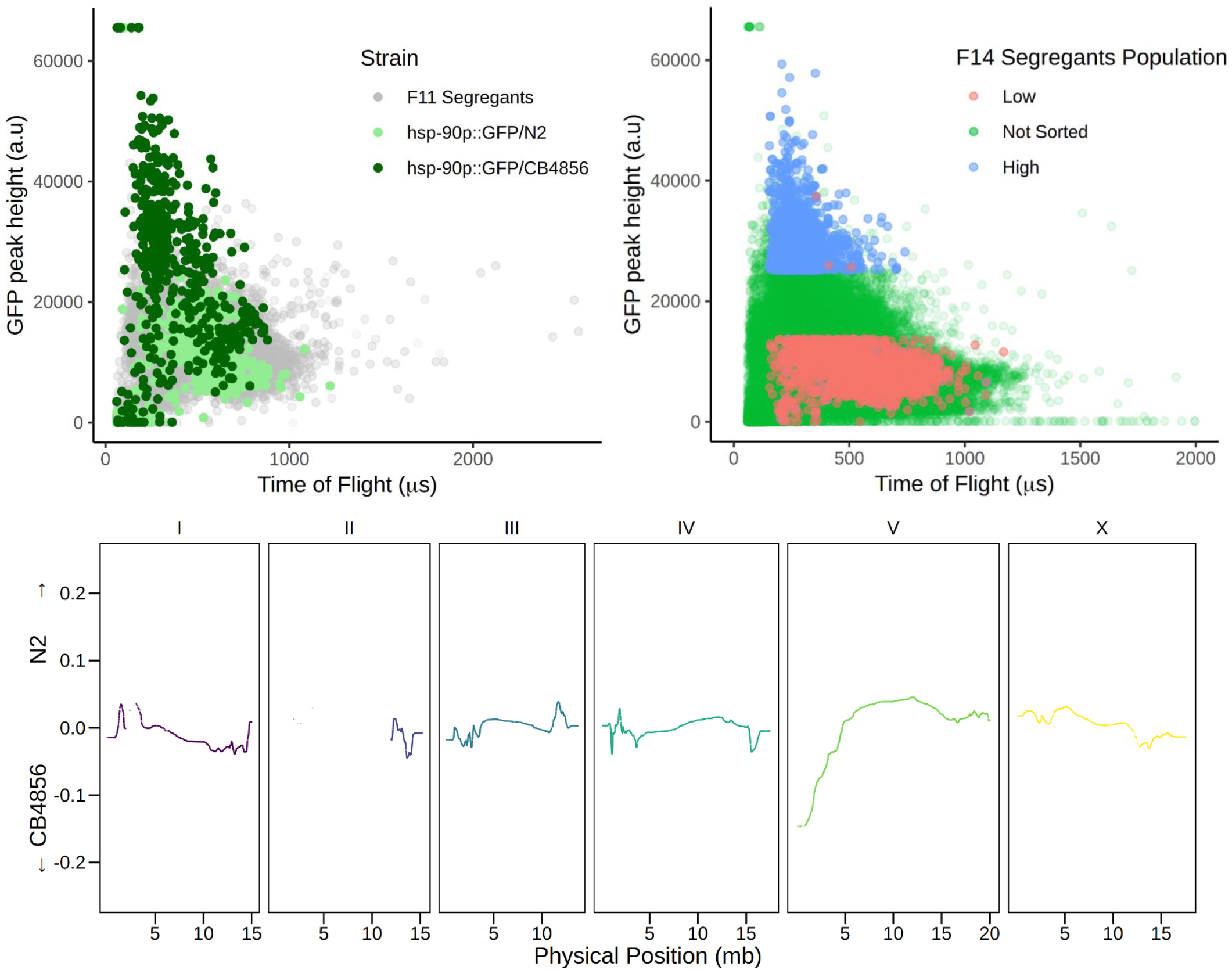
X-QTL mapping for *hsp-90* expression levels. **(A)** Sorting of the parental X-QTL strains and F11 segregants (Left), and F_14_ N2 × CB4856 segregant pool carrying the *hsp-90*::GFP transcriptional reporter (Right). **(B)** Differences in allele frequencies between the ‘High’ and ‘Low’ GFP fluorescence sorted pools. The ‘High’ GFP expression pool is enriched for CB4856 at the QTL locus in Chr. V.

**Supplementary Fig. 8.**
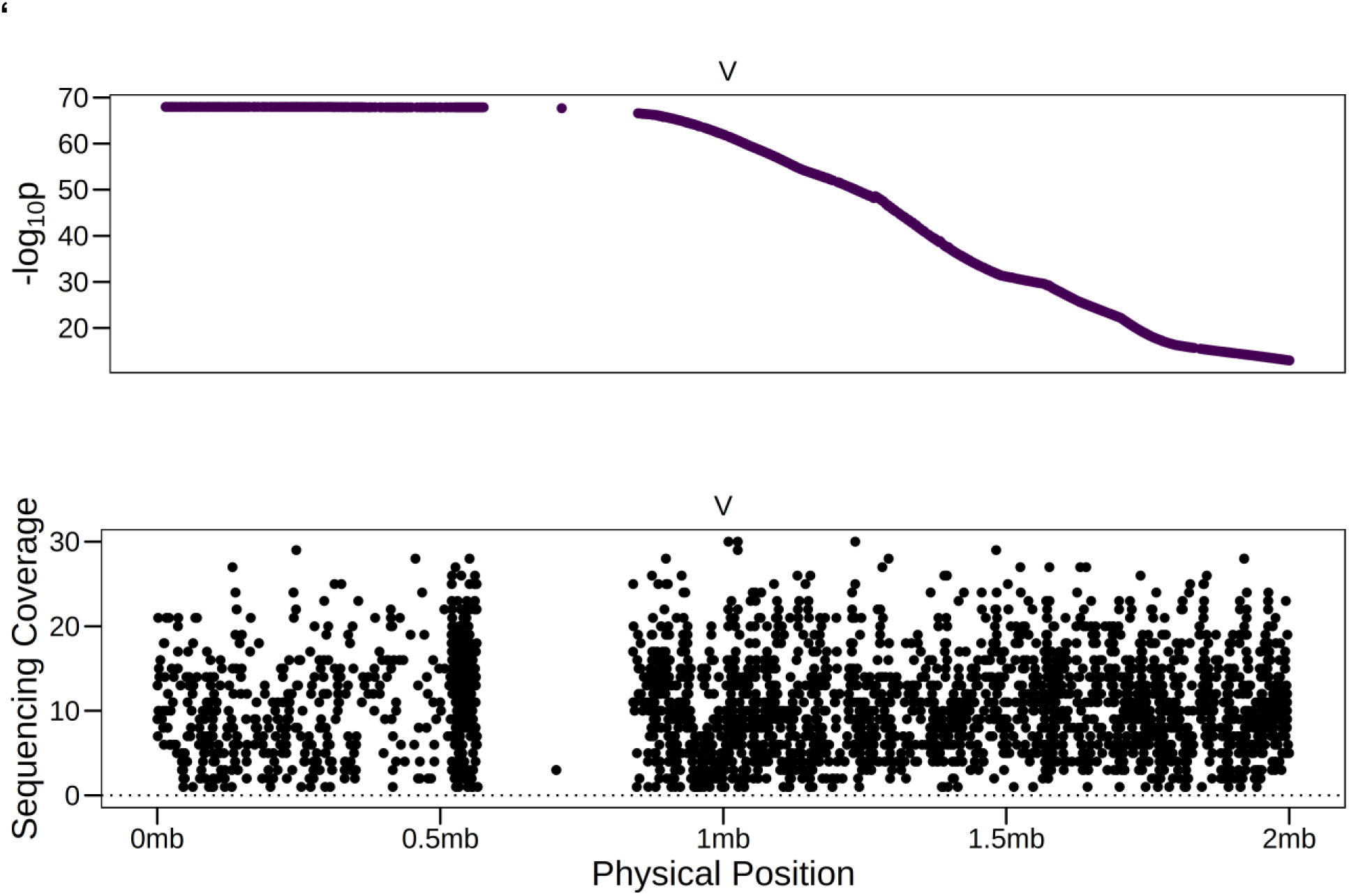
The CB4856 *fog-2(q71) hsp-90*::GFP strain contains a large de novo deletion. (Top) *p-value* for association with GFP signal on the left end of Chr. V. SNVs that are fixed in the population (90% or above) are removed from the analysis. (Bottom) The coverage at SNVs in a strain showing upregulated GFP expression reveals a large region that is depleted, indicating a deletion.

**Supplementary Fig. 9.**
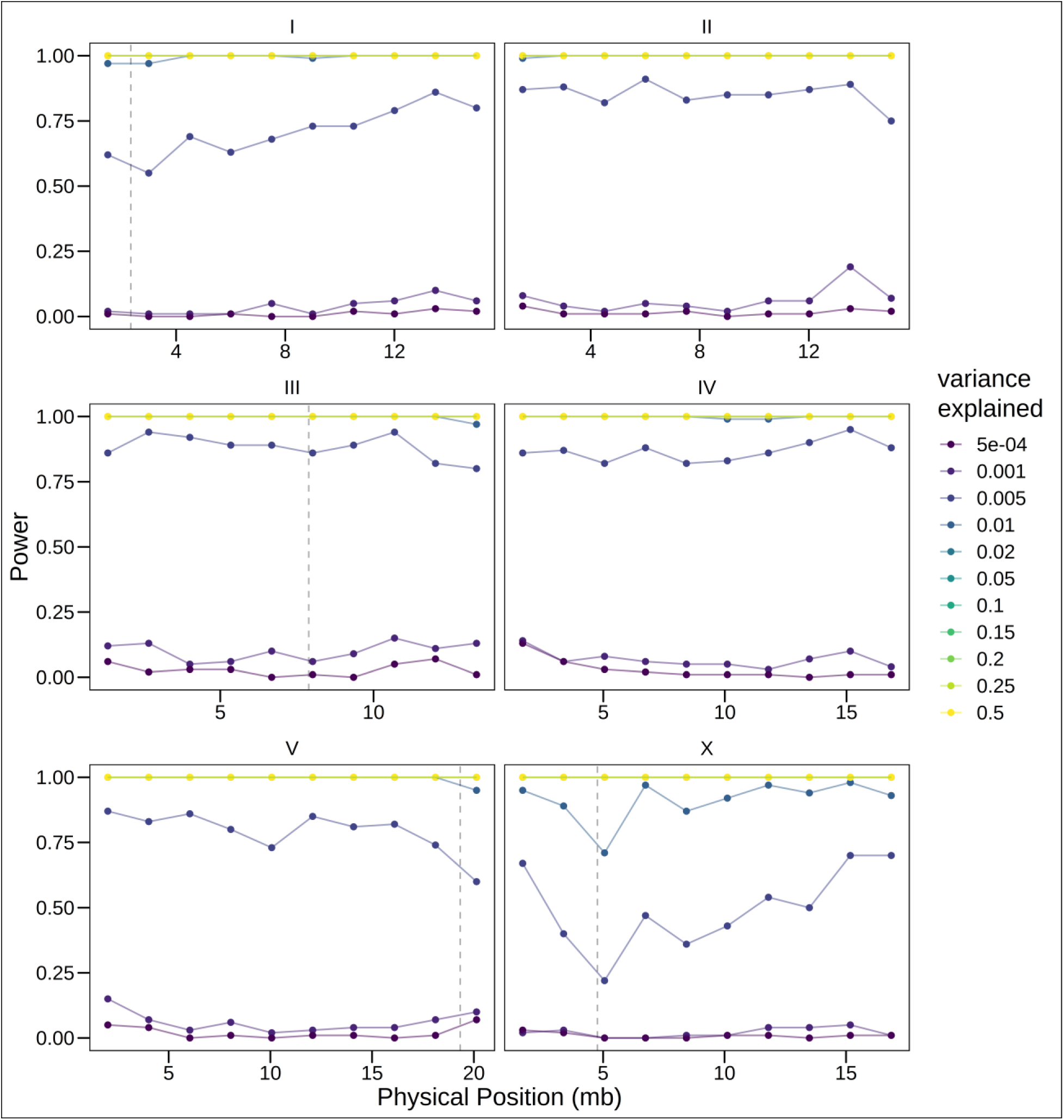
Baseline changes in allele frequencies due to large fitness effects do not prevent X-QTL mapping. Simulations were carried out as described in Supplementary Figure S2, except that we simulated the effect of *peel-1/zeel-1* on Chr I, as well as three loci conferring a fitness advantage on chromosomes III, V, and X as observed in our data (dashed lines). In our simulations, only 20% of the males participate in mating, and we simulated fitness affecting loci as modifiers of the probability of a male to be selected for mating. The specific parameters for each locus were determined by identifying those that resulted in selection strength similar to observed effects.

**Supplementary Fig. 10.**
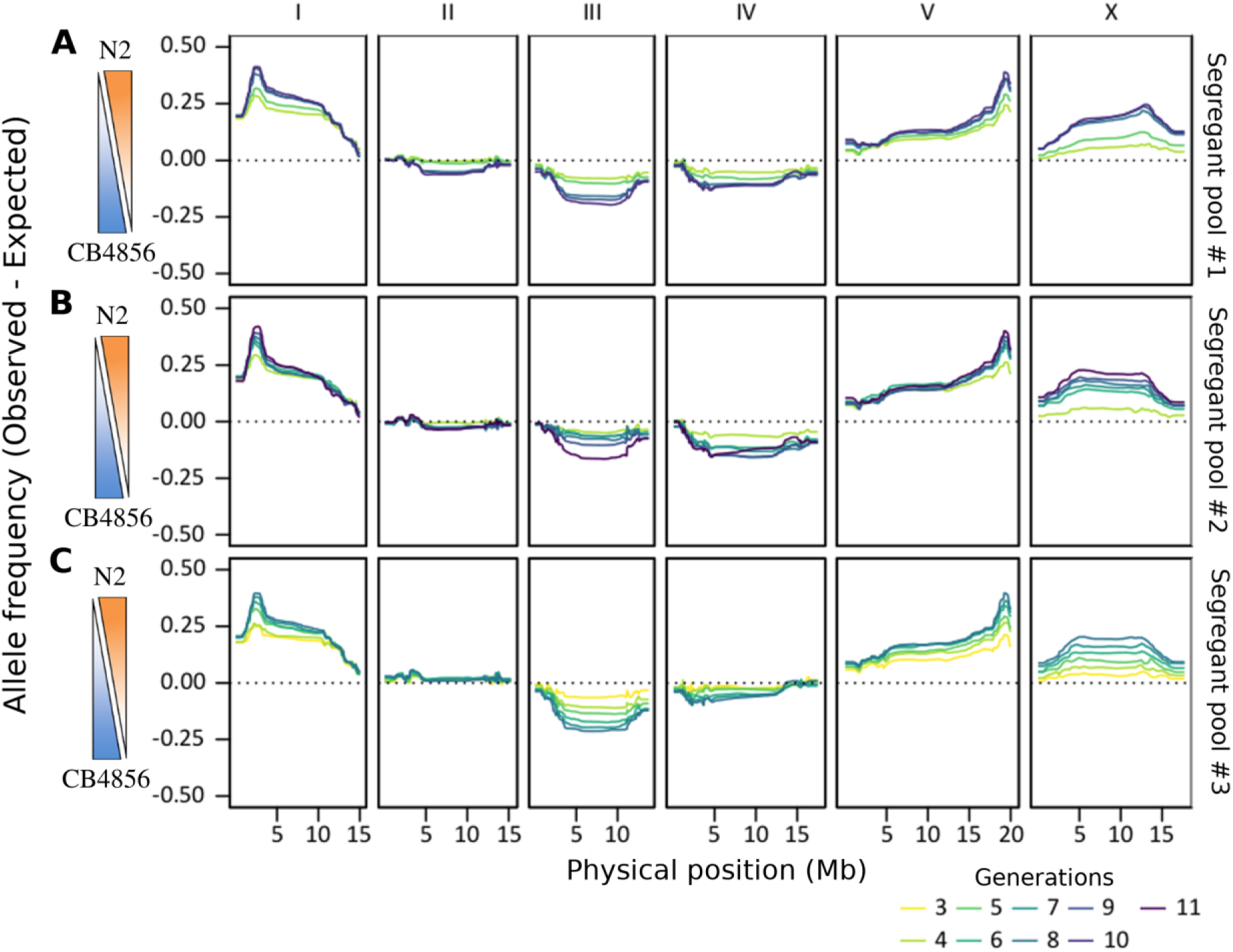
High reproducibility of changes in allele frequency in X-\QTL segregant populations. **(A-C)** Changes in allele frequencies across multiple generations (ranging from F_3_-F_11_) in a N2 × CB4856 cross. Allele frequencies correspond to the difference between the observed and the expected values. Three biological repeats are shown. Each of these populations originates from an independent cross of parental strains and they were grown months apart.

**Supplementary Fig. 11.**
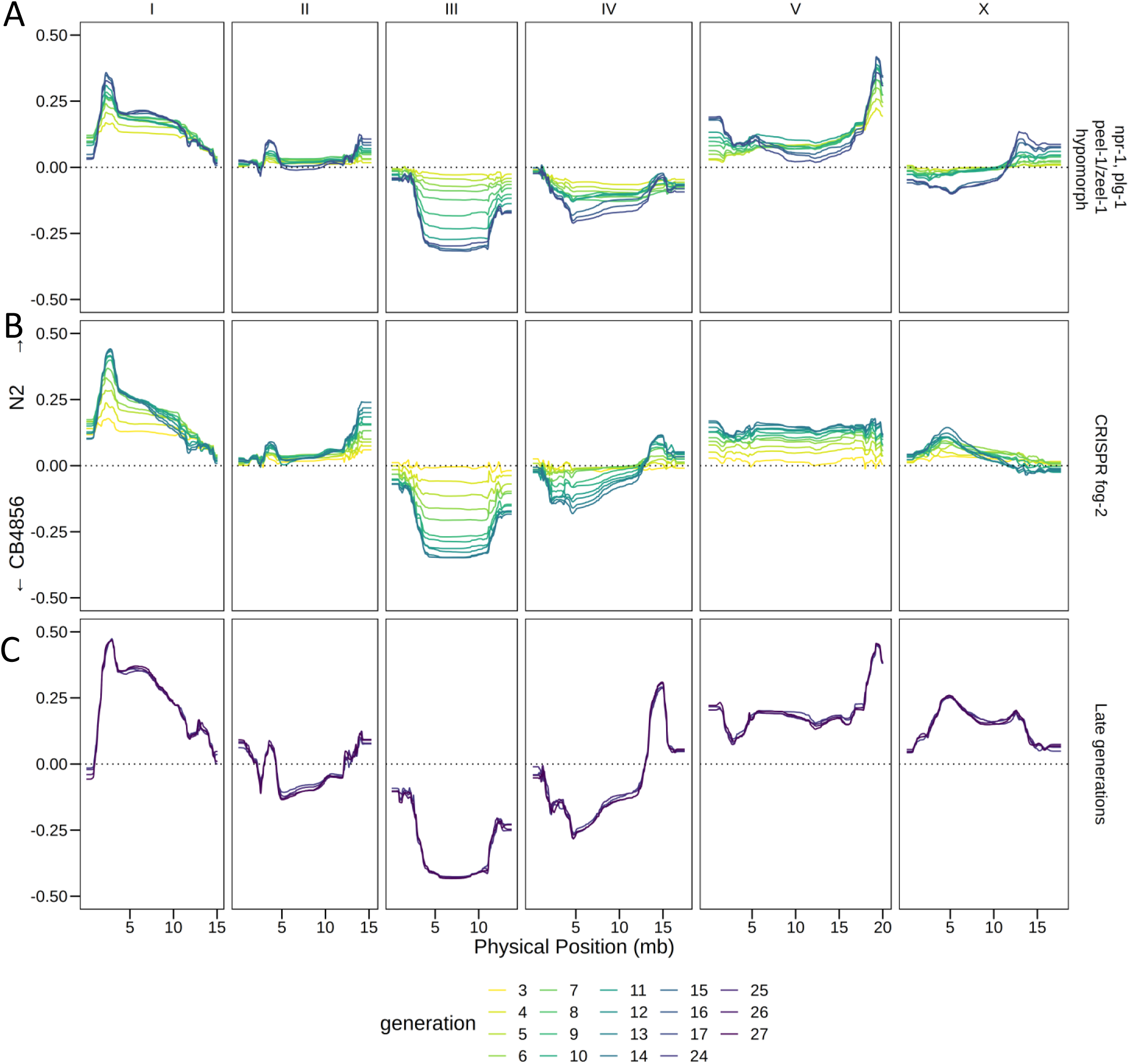
Additional X-QTL segregant pools. **(A)** We crossed the parental CB4856 *fog-2(q71)* strain to a modified N2 *fog-2(q71)* strain carrying a hypomorphic *peel-1/zeel-1 (ttTi12715)* allele (Chr. I, introgressed from QX1430), a functional *plg-1* allele (Chr. III, introgressed from CB5203) and the WT allele of *npr-1*. Drive towards N2 in the *peel-1/zeel-1* locus is weaker and selection towards N2 *npr-1* is completely abolished. No changes were observed in Chr. III indicating that variation in *plg-1* does not underlie the selection in favor of CB4856 alleles (compare (A) to Supplementary Figure 10). **(B)** Dynamics of an X-QTL segregant pool generated by replacing the “introgression CB4856 *fog-2(q71)* parental strain” with a CB4856 *fog-2* strain generated by CRISPR/Cas9 targeted allele replacement. Most large fitness peaks are reproduced (compare (B) to Supplementary Figure 10) with the exception of a locus linked to fog-2 (right end of Chr. V) and a secondary peak in Chr X. **(C)** allele frequencies of advanced X-QTL generations (F_24_-F_27_). Two loci of antagonizing effects on fitness are identified on Chr. IV selecting for CB4856 (left arm of the chromosome) and N2 (right arm).

## Supplementary Tables

**Supplementary Table 1**. Details of RNAi screen to identify the transcriptional regulator of *hsp-90*.

**Supplementary Table 2**. *C. elegans* strains used in this study

**Supplementary Table 3**. Illumina short-read sequencing runs

